# Fast and background-suppressed DNA-PAINT

**DOI:** 10.64898/2026.11.10.765791

**Authors:** Tingting Sun, Shuting Liu, Xi Zhang, Huang Tang, Boyang Hua, Xing-Hua Xia

**Affiliations:** State Key Laboratory of Analytical Chemistry for Life Science, School of Chemistry, Nanjing University, Nanjing 210023, China; Department of Oncology, Nanjing Drum Tower Hospital, Nanjing University, Nanjing 210008, China

**Keywords:** photoactivatable fluorescent probes, photoactivation, diffusion, excitation (PhADE), spinning disk confocal (SDC) microscopy, super-resolution imaging

## Abstract

The speed and resolution of DNA-PAINT imaging is partly limited by the background fluorescence at high imager-strand concentrations. Here, we combine the photoactivatable fluorophore-quencher-DNA probes with the photoactivation, diffusion, and excitation strategy on a spinning disk confocal microscope, decoupling background fluorescence from the imager-strand concentration. This method yields >7-fold faster DNA-PAINT imaging in cells, a ∼15 nm localization precision in hydrogels, without using specialized or speed-optimized imager strands.

---

Single-molecule localization microscopy (SMLM) represents a broad collection of powerful tools that have become essential for modern-day biological research.^1^ In SMLM, to resolve individual molecules beyond the diffraction limit, only a sparse subset of fluorophores is switched on at any given time. For example, DNA point accumulation for imaging in nanoscale topography (DNA-PAINT)^2^ utilizes the transient binding and dissociation of imager strands to achieve the stochastic switching of fluorophores. However, imposed by this signal sparsity, the trade-off for high spatial resolution in SMLM is the long image-acquisition time. Although numerous algorithms have been developed to localize densely spaced single molecules,^3, 4^ implementing the high-density scenario in DNA-PAINT still faces several practical limitations. For example, although a higher image-acquisition speed can be achieved via a higher binding rate *k*_*on*_, such as those through optimizing sequence design and buffer conditions,^5^ or through tandem binding sites,^6^ the number of such optimized imager strands remain limited. On the other hand, increasing the imager-strand concentration would be a general and facile route to achieve the same purpose, but the resulting higher background fluorescence could substantially deteriorate the localization resolution and precision. Recently, DNA-PAINT has been applied in hydrogel-imprinted and cleared samples.^7^ However, the dense hydrogel matrices render hybridization inefficient and therefore necessitate the use of higher imager-strand concentrations, further exacerbating the issue of background fluorescence. In this context, FRET DNA-PAINT,^8^ PA DNA-PAINT,^9^ TDI DNA-PAINT,^10^ and fluorogenic DNA-PAINT^11, 12^ offer a compelling solution to the issue by suppressing background fluorescence, although these methods are constrained in their probe designs. In this study, we achieved a >7-fold increase in imaging speed in DNA-PAINT by increasing the imager-strand concentration while suppressing the background fluorescence via the use of a recently developed fluorophore-quencher-DNA (FQD) probe together with the photoactivation, diffusion, and excitation (PhADE) strategy. We anticipate that the FQD-based DNA-PAINT approach could be broadly adopted for super-resolution imaging in diverse biological contexts.

To achieve fast and background-suppressed DNA-PAINT, we combined the unimolecular FQD (FQD1) probes with PhADE, and implemented the method on a confocal microscope.^13^ In this approach, the FQD1594-labeled imager strands are maintained in a non-fluorescent “off” state due to efficient quenching. Upon 405-nm illumination, a fraction of the FQD probes in the excitation volume is photoactivated; however, only those that are transiently bound to the immobilized docking strands remain observable during the subsequent 561-nm excitation, while the unbound photoactivated probes rapidly diffuse out of the confocal detection volume (Figs. 1a,b). This diffusion-assisted background suppression allows the use of high imager-strand concentrations without sacrificing the signal-to-background ratio (SBR), thereby potentially increasing the number of binding events per frame and accelerating image acquisition.

**Fig. 1.**
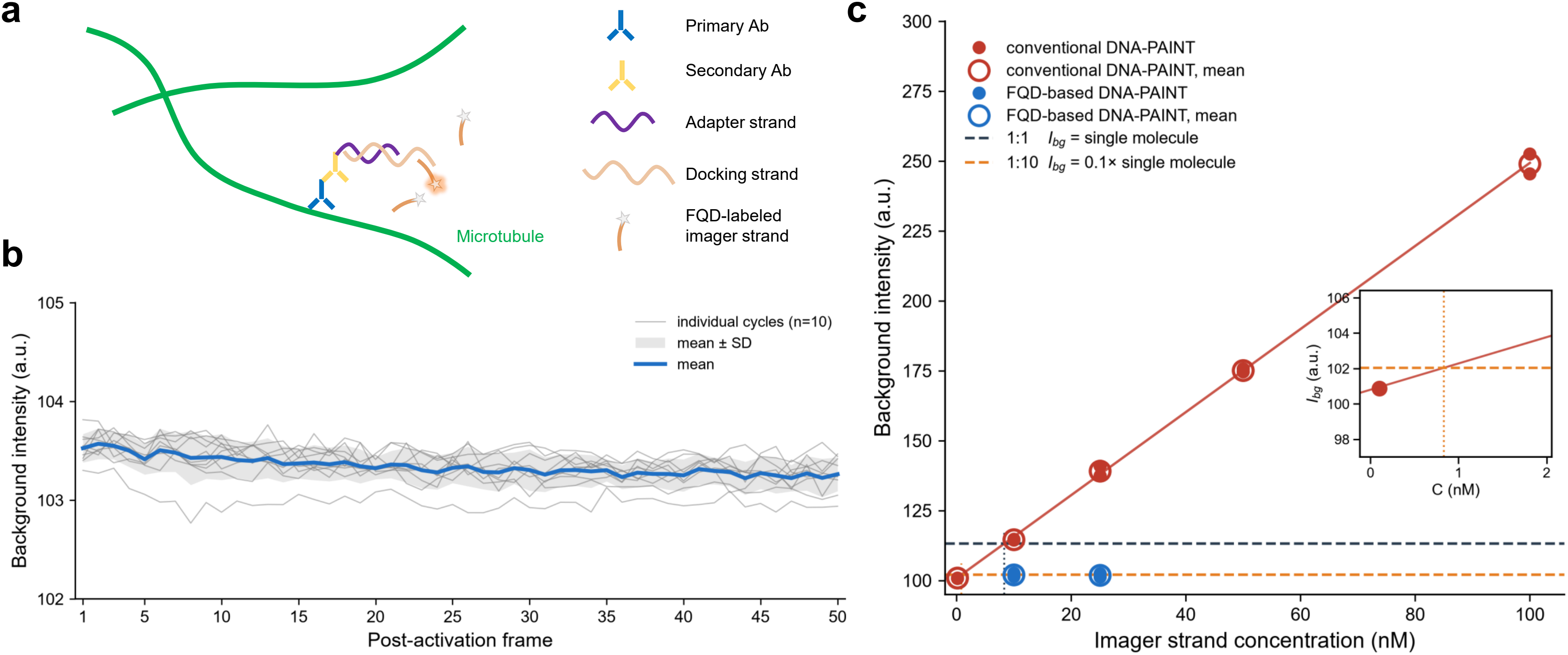
Background suppression in DNA-PAINT by SDC PhADE. **a**, Workflow of FQD-based SDC PhADE DNA-PAINT. **b**, Background fluorescence vs. post-activation frame for FQD-based SDC PhADE DNA-PAINT at a 10 nM imager-strand concentration. **c**, Background fluorescence vs. imager-strand concentration for conventional and FQD-based SDC PhADE DNA-PAINT. I_bg_ = 1.49 × C + 100.8, R^2^ = 0.998. Photoactivation of 1 frame followed by diffusion and excitation of 50 frames were performed in each SDC PhADE cycle.

To test if confocal PhADE can achieve nanometer-scale localization resolution and precision from single fluorophores under high concentrations of imager strands, we chose to observe DNA hybridization events in hydrogels,^13^ as hydrogel-embedded samples typically require high imager-strand concentrations to achieve efficient hybridization kinetics.^7^ Specifically, we immobilized the Acrydite-AF488-docking strands in the hydrogel matrices, and added in the complementary FQD1594-imager strands through diffusion.^13^ Single-molecule spots were subsequently observed in the confocal PhADE images, whose centroid localization was fitted and clustered in a 2D histogram. We determined the single-molecule localization resolution and precision to be about 35 nm and 15 nm, respectively, in the xy-direction (full width at half maximum, FWHM = ∼2.35σ),^13^ similar to those obtained by related DNA-PAINT techniques.^7, 14^ Moreover, the localization resolution did not deteriorate at higher concentrations of imager strands (up to 200 nM), demonstrating the effectiveness of background suppression by confocal PhADE.^13^ These results confirm that the combination of FQD probes and confocal PhADE preserves the single-molecule localization resolution and precision even under high concentrations of imager strands.

To test if the FQD-based confocal PhADE method can be applied to DNA-PAINT imaging in cells, we first characterized the concentration-dependent background fluorescence of different DNA-PAINT methods. Here, we switched to spinning disk confocal (SDC) microscope for a larger field of view and higher frame rates, which are essential for super-resolution cellular imaging. For conventional DNA-PAINT, the background intensity was observed to increase linearly with the imager-strand concentration (Fig. 1c, from 100 pM to 100 nM). Based on the single-molecule intensity measured at 100 pM (median = 12.4 a.u., n = 100), the background intensity would reach 10% of the single-molecule signal at approximately 0.8 nM, and the SBR would further drop to 1 at approximately 8 nM (Fig. 1c). These results underscore the intrinsic concentration limit of conventional DNA-PAINT, i.e., increasing the imager-strand concentration to accelerate binding inevitably elevates the background, degrading localization precision and making single-molecule detection increasingly difficult. In contrast, when the FQD1594-imager strands were used under the SDC PhADE configuration, the background remained around 10% of the single-molecule signal even at a concentration of 25 nM (Fig. 1c), demonstrating that SDC PhADE effectively decouples background fluorescence the from the imager-strand concentration.

We then compared the imaging performance of conventional and FQD-based DNA-PAINT on microtubule structures in fixed cells. With conventional DNA-PAINT at 200 pM, the number of single-molecule spots per frame was low, and even after accumulating around 3,000 frames, the microtubule structure remained poorly defined (resolution = 82 nm,^15^ Fig. 2a). In stark contrast, FQD-based DNA-PAINT at 5 nM yielded a markedly higher spot density per frame while maintaining a low background, and only about 500 frames were sufficient to reconstruct a clear microtubule structure (resolution = 88 nm,^15^ Fig. 2b). The >7-fold increase in imaging speed achieved here is attributed to the higher imager-strand concentrations enabled by SDC PhADE. Notably, the SDC PhADE approach substantially improved both the imaging area and the temporal resolution compared with our previous confocal PhADE implementation,^13^ making it practical for routine super-resolution cellular imaging.

**Fig. 2.**
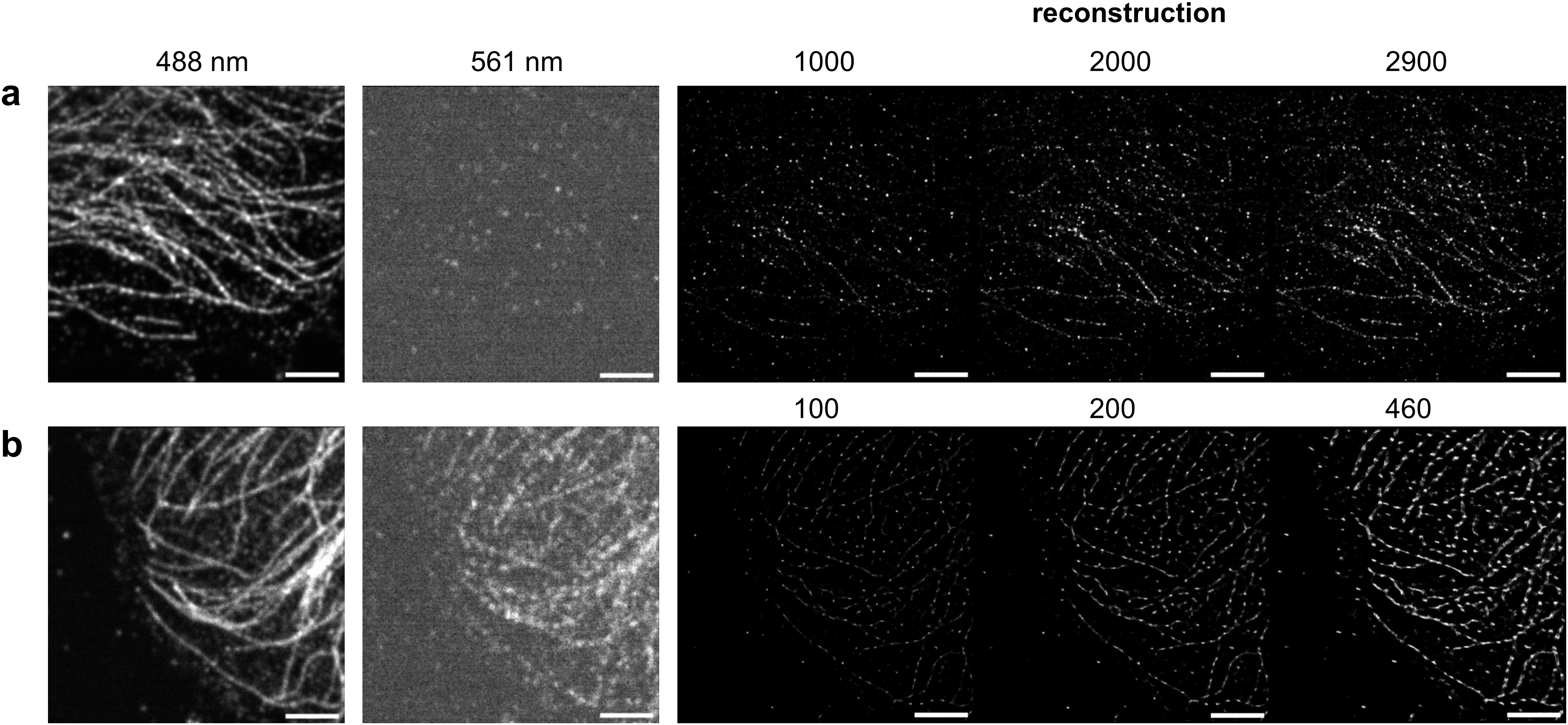
Fast and background-suppressed DNA-PAINT imaging of microtubules. **a**, Conventional DNA-PAINT at a 200 pM imager-strand concentration at various frame numbers. Accumulation of 2900 frames, containing ∼150000 total localizations, yield a poorly defined microtubule structure. The image size was 16.64 × 16.64 µm^2^. **b**, FQD-based SDC PhADE DNA-PAINT at a 5 nM imager-strand concentration at various frame numbers. Accumulation of 460 frames, containing ∼181000 total localizations, yield a clear microtubule structure. The image size was 16.64 × 16.64 µm^2^. Scale bars, 3 µm. Photoactivation of 1 frame followed by diffusion and excitation of 5 frames were performed in each SDC PhADE cycle.

A key advantage of our approach lies in its simplicity and accessibility. The FQD1 probes are minimally 2 nt longer than conventional imager strands, flexible with fluorophores, and can be ordered commercially without the need for custom synthesis.^13^ The only additional requirement is a standard 405-nm laser line for photoactivation, which is available on most commercial confocal microscopes. In contrast, other background-suppression strategies for DNA-PAINT often require specialized probes. For example, FRET DNA-PAINT requires dual-labeled donor–acceptor pairs and suffers from photobleaching of the acceptor-labeled docking strand, whereas PA DNA-PAINT works with a limited choice of fluorophores that can be reductively caged. In addition, fluorogenic DNA-PAINT typically requires substantially longer imager strands (15 nt), which have a higher propensity for secondary structures.^11^

The compatibility with SDC microscopy further enhances the practicality of the method. SDC microscopy extends DNA-PAINT to deep volumetric imaging,^14^ but is typically more sensitive to background fluorescence than total internal reflection fluorescence (TIRF) microscopy. We anticipate that the combination of FQD probes, SDC microscopy, and PhADE will be broadly applicable to a wide range of samples, including fixed cells, tissue sections, and hydrogel-embedded or cleared specimens, where high imager-strand concentrations are essential for efficient hybridization. Future work could explore the extension of this method to multicolor imaging and to live-cell super-resolution applications, where the rapid acquisition and low background would be highly beneficial.

In summary, we have established a fast, background-suppressed DNA-PAINT method that leverages commercially available FQD probes and a simple PhADE strategy on a standard SDC microscope. The method achieves >7-fold speed improvement, preserves nanometer localization precision, and is compatible with hydrogel-embedded samples. We expect that this approach will facilitate the broader adoption of DNA-PAINT for deep volumetric super-resolution imaging in diverse biological contexts.

## Supporting information

Experimental section

## Acknowledgments

We thank all members of the Boyang Hua group for helpful discussion. Besides, we acknowledge and thank Dr. Yi Shi and Dr. Weikang Wang at the School of Chemistry and Molecular Engineering, East China Normal University for providing the confocal microscope used in this study and related technical support. This work was supported by the National Natural Science Foundation of China (Nos. 22404079, U24A20503), the Fundamental Research Funds for the Central Universities 2024300410, and the State Key Laboratory of Analytical Chemistry for Life Science, Nanjing University 5431ZZXM2403.

## Author contributions

**T.S**.: Methodology, Investigation, Formal analysis, Writing–Original Draft, Writing–Review & Editing;

**S.L**.: Methodology, Investigation, Formal analysis, Writing–Original Draft, Writing–Review & Editing;

**X.Z**.: Investigation;

**H.T**.: Investigation;

**B.H**.: Conceptualization, Methodology, Writing–Original Draft, Writing–Review & Editing, Supervision, Funding acquisition;

**X.X**.: Supervision, Funding acquisition

