## Supplementary material for "Fast and background-suppressed DNA-PAINT": Experimental section

**Materials**

Sample chambers were ordered from Cellvis (C18SB-1.5H and C8SB-1.5H). HEPES solution (Sigma-Aldrich, 83264), Tris-HCl (pH 7.4, Beyotime, ST786), Tris-HCl (pH 8.0, Beyotime, ST786), PBS (pH 7.4, KeyGen, KGL2206-500), potassium chloride (Aladdin, P112133), magnesium chloride (Sigma-Aldrich, 442615), EDTA (ThermoFisher, AM9260G), glucose (Sigma-Aldrich, G7528), (+-)-6-hydroxy-2,5,7,8-tetra-methylchromane-2-carboxylic acid (Trolox, Sigma-Aldrich, 238813), glucose oxidase (Sigma-Aldrich, G2133), catalase (Sigma-Aldrich, C9322), Alconox (Sigma-Aldrich, Z742914), potassium hydroxide (Sigma-Aldrich, 484016), triethoxyvinylsilane (Sigma-Aldrich, 679275), ethanol (Aladdin, E111993), acetic acid (Aladdin, A298827), 40% acrylamide-bisacrylamide (19:1) (Beyotime, ST004B), APS (Sigma-Aldrich, A3678), TEMED (Sigma-Aldrich, T7024), Tween 20 (Sigma-Aldrich, T8787), HEPES-free, high-glucose Dulbecco's modified Eagle's medium (DMEM, Sangon, E600002-0500), 10% fetal bovine serum (FBS, Vazyme, F101-01), penicillin-streptomycin (MCE, HY-K1006), 4% paraformaldehyde fixative solution (Beyotime, P0099-100ml), Triton X-100 (Sigma-Aldrich, P9416-100ML), immunostaining blocking buffer (Beyotime, P0102), immunostaining primary antibody diluent (Beyotime, P0103), immunostaining (non-fluorescent) secondary antibody diluent (Beyotime, P0110), NeutrAvidin (Thermo Scientific, 31000), 20X SSC (Beyotime, ST463-500ml), yeast tRNA (Beyotime, R0040-0.5ml), nuclease-free water (Beyotime, ST876-500ml), Beta Tubulin Rabbit recombinant antibody (Proteintech, 80713-1-RR), biotin-conjugated Goat anti-Rabbit IgG (H+L) (Proteintech, SA00004-2).

**Buffers**

Buffer A was 25 mM HEPES, pH 7.6, 150 mM KCl, ultrapure H_2_O.

Buffer C was 5 mM Tris-HCl pH 8, 75 mM MgCl_2_, 0.05% Tween-20, 1 mM EDTA, 0.8% glucose, 4 mM Trolox, ultrapure H_2_O, and 1X Gloxy was added before imaging.^1^

Buffer F was 1X PBS, 360 mM NaCl, 0.8% glucose, 4 mM Trolox, ultrapure H_2_O, and 1X Gloxy was added before imaging.^1^

**Oligonucleotide preparation**

AF594- and FQD_1_594-imager strands, Acrydite-AF488-docking strand (10 nt), Acrydite-docking strand (9 nt), and AF488-docking strand-biotin were purchased from Sangon Biotech.

To generate Acrydite-AF488-docking strand-biotin (9 nt), Acrydite-docking strand (9 nt) was hybridized with AF488-docking strand-biotin. DNA hybridization was carried out by heating and slow cooling samples from 95 °C to 30 °C in approximately 10 min in Buffer A. All oligo samples were stored at -20 °C.

**Imaging surface preparation**

For the triethoxyvinylsilane-modified imaging chambers, they were prepared according to the protocol described previously.^2^ Briefly, the imaging chambers were sequentially sonicated in Alconox solution, ultrapure H_2_O, and KOH solution. The imaging chambers were then thoroughly rinsed with ultrapure H_2_O and dried with nitrogen gas. The imaging chambers were then subjected to an oxygen plasma treatment, followed by an overnight silane treatment. Finally, the imaging chambers were thoroughly rinsed with ethanol and dried with nitrogen gas.

**Confocal microscopy**

The DNA hybridization experiments were carried out in the fluorescence recovery after photobleaching (FRAP) mode on a Leica Stellaris 8 STED microscope as described previously,^2^ which is equipped with a 405-nm laser, a white light laser (WLL) source, and an HC PL APO CS2 100× oil-immersion objective (NA = 1.4). Briefly, AF594 was excited with the 580-nm laser from the WLL source, and the fluorescence was collected in the photon-counting mode (gate time 0.3–6.0 ns) between 600–700 nm (for AF594) using a HyD detector. The scanning speed was 102.38 μs/pixel, and the pixel size was 51.53 × 51.53 nm^2^. The z-depth, image size, and laser power were specified in the figure captions for each experiment.

Based on the pixel dwelltime of 102.38 μs and pixel size of 51.53 nm on the microscope, the calculated 405-nm photoactivation time per molecule (~350 × ~350 nm^2^) was 5.01 ms per frame. The effective diffusion time in a PhADE cycle was the time interval between two consecutive confocal scans of the same molecule.

**Confocal PhADE** **imaging of individual DNA hybridization events**

The hydrogel samples containing the Acrydite-AF488-docking strand (10 nt) were prepared on triethoxyvinylsilane-functionalized imaging chambers according to the protocol described previously.^2^ All the preparation steps were performed on ice until polymerization. The imaging chambers were then washed three times with 300 μL of 1X PBS buffer, and the FQD_1_594-imager strands (20, 50, 100, 200 nM) in Buffer C were added to the hydrogel for 1 h before 1X Gloxy was added. Confocal PhADE was carried out on a confocal microscope (see confocal microscopy), which includes the photoactivation (405 nm, equivalent to a single-molecule activation time of 5.01 ms per frame) and excitation (594 nm, approximately 520 ms per frame) steps in each confocal PhADE cycle. For each experiment, the detailed imaging parameters are described in the corresponding figure captions.

**Cell culture**

The complete medium was prepared by adding 50 mL FBS and 5 mL penicillin-streptomycin to 500 mL DMEM (high glucose, without HEPES). HeLa cells were cultured in a 37 °C incubator with 5% CO_2_.

**Fixation and immunofluorescence**

The culture medium was aspirated, and the cells were washed twice rapidly with 500 μL 1X PBS per well. The cells were fixed with 200 μL of 4% paraformaldehyde at room temperature for 15 min without shaking. After fixation, the fixative was aspirated, and the cells were washed three times with 300 μL 1X PBS for 5 min each. The cells were permeabilized with 200 μL of 0.2% Triton X-100 in 1X PBS at room temperature for 10 min without shaking. After permeabilization, the permeabilization solution was aspirated, and the cells were washed three times with 300 μL 1X PBS for 5 min each. The cells were blocked with 200 μL of immunostaining blocking buffer for 1 h. For primary antibody staining, 4 μL Beta Tubulin Rabbit recombinant antibody was diluted in 796 μL immunostaining primary antibody diluent (1:200). After aspirating the blocking buffer, 100 μL of the primary antibody solution was added to each well, and the 8-well plate was incubated at 4 °C overnight on a shaker. The primary antibody solution was aspirated, and the cells were washed three times with 300 μL 1X PBS for 5 min each. For secondary antibody staining, 8 μL biotin-conjugated Goat anti-Rabbit IgG (H+L) was diluted in 792 μL immunostaining secondary antibody diluent (1:100). 100 μL of the secondary antibody solution was added to each well, and the 8-well plate was incubated for 1 h. The secondary antibody solution was aspirated, and the cells were washed three times with 300 μL 1X PBS for 5 min each. All steps were performed at room temperature on a shaker at 50 rpm unless otherwise specified.

For NeutrAvidin (NA) incubation, NA (1 μg/μL) was diluted 1:1000 in immunostaining secondary antibody diluent (0.8 μL NA in 800 μL diluent). 200 μL of the NA working solution was added to each well, and the 8-well plate was incubated for 30 min. The NA working solution was aspirated, and the cells were washed three times with 300 μL 1X PBS for 5 min each. The docking strand diluent (1 mL) was prepared by mixing 100 μL 20X SSC, 100 μL 1% Tween 20, and 20 μL yeast tRNA in 780 μL nuclease-free water. The docking strand working solution (10 nM) was prepared by adding 8 μL of 1 μM docking strand to 792 μL docking strand diluent. 100 μL of the docking strand working solution was added to each well, and the 8-well plate was incubated for 30 min. The docking strand working solution was aspirated, and the cells were washed three times with 300 μL 1X PBS for 5 min each. All steps were performed at room temperature on a shaker at 50 rpm under light-protected conditions unless otherwise specified.

**Spinning disk confocal microscopy**

The SDC PhADE experiments were carried out in the spinning disk confocal mode on a Cell Xpanse live-cell wide-field all-time super-resolution microscope with a 405-nm laser, a 488-nm laser, a 561-nm laser, and a 100× oil-immersion objective (NA = 1.5). Briefly, AF488 and AF594 were excited with the 488-nm and 561-nm lasers, respectively, and the fluorescence was collected between 515–535 nm (for AF488) and 592–626 nm (for AF594) using a sCMOS detector. The pixel size was 65 × 65 nm^2^. The image size was specified in the figure captions for each experiment.

**SDC PhADE DNA-PAINT imaging of microtubules**

The AF594- and FQD_1_594-imager strands in Buffer F at the specified concentrations were added to the labeled cells with 1X Gloxy. SDC PhADE was carried out on a SDC microscope (see spinning disk confocal microscopy), which includes the photoactivation (405 nm, 100 ms per frame) and excitation (594 nm, 200 ms per frame) steps in each SDC PhADE cycle. For each experiment, the detailed imaging parameters are described in the corresponding figure captions.

**Data analysis**

Single-molecule centroid localization analysis for DNA hybridization experiments and DNA-PAINT reconstruction were performed using ThunderSTORM in ImageJ and custom-made scripts in MatLab. Specifically, drift correction with the fiducial marker (DNA hybridization) or cross-correlation (DNA-PAINT) method was applied to the raw localization data to generate the drift-corrected 2D histograms. The FWHM values were calculated by 2D Gaussian fitting for individual localization event clusters implemented in MatLab. The DNA-PAINT image resolution was estimated with decorrelation analysis in ImageJ.^3^

**Table 1** The Oligo list.

| **Names** | **Sequences (5’ to 3’) and modifications** |
| --- | --- |
| **imager strands** | **FQD_1_594-imager strand**  BHQ2/T/iPCLink/T/iAF594dT/TTCTAGATGTAT  **AF594-imager strand**  AF594-TTTTTCTAGATGTAT |
| **docking strands** | **Acrydite-AF488-docking strand (10 nt)**  Acrydite-ATCCC/iAF488dT/ATCTTATACATCTAG  **Acrydite-docking strand (9 nt)**  Acrydite-CTTTTCTTCTTTTCTTTCTTTTCCTCCTCTACATCTAG  **AF488-docking strand-biotin**  AAGAAAGAAAAGAAGAAAAGT/iAF488dT/TT-biotin |

***References***

(1) Rasnik, I.; McKinney, S. A.; Ha, T. Nonblinking and long-lasting single-molecule fluorescence imaging. Nat Methods 2006, 3 (11), 891–893.

(2) Tang, H.; Liu, S.; Zhang, X.; Xia, X.-H.; Hua, B. Generalized photoactivatable fluorescent probes enable background-suppressed 3D single-molecule imaging in hydrogels. Research Square 2026, rs–9070027.

(3) Descloux, A.; Grußmayer, K. S.; Radenovic, A. Parameter-free image resolution estimation based on decorrelation analysis. Nature Methods 2019, 16 (9), 918–924.
